# Dual effects of nicotinamide on aging-related arrhythmia: protective at low dose, proarrhythmic at high doses

**DOI:** 10.1101/2025.09.22.677785

**Authors:** Liangyu Hu, Yun Hu, Xi Qi, Alexia van Rinsum, Jaap Keijer, Deli Zhang

## Abstract

**Background:** Cardiac arrhythmia and dysfunction progressively increase with age, both of which are closely associated with a decline in NAD^+^. Supplementation with nicotinamide (NAM), a critical NAD^+^ precursor, has shown protection in experimental models of aging-related cardiac diseases, including atrial fibrillation (AF) and heart failure, especially heart failure with preserved ejection fraction. With the potential rise of NAM in treatment of various cardiometabolic diseases, its dose-dependent cardiac-specific adverse effects and underlying mechanisms warrant investigation.

**Methods:** The effects of NAM on aging-related arrhythmia and cardiac dysfunction were assessed using *ex vivo* Langendorff mouse hearts and adult *Drosophila* heart preparations. Different doses of NAM (10-100 mM) were tested for their impact on the contractility of HL-1 cardiomyocytes, lifespan and cardiac function of *Drosophila*, as well as arrhythmia susceptibility of e*x vivo* mouse hearts. Acetylation of sarcoplasmic/endoplasmic reticulum Ca²⁺ ATPase 2a (SERCA2a) was measured by immunoprecipitation followed by Western blotting.

**Results:** Acute perfusion with 10 mM NAM had limited influence on aging-related AF susceptibility in *ex vivo* mouse hearts. Short-term dietary intervention with 10 mM NAM in late-life protected against aging-induced cardiac arrhythmia and contractile dysfunction exclusively in male *Drosophila*. In contrast, life-long exposure or NAM concentrations above 20 mM led to dose-dependent adverse cardiac effects, including impaired contractility in HL-1 cardiomyocyte and shortened lifespan in *Drosophila*, with increased arrhythmia observed in both models. In *ex vivo* mouse hearts, 100 mM NAM increased SERCA2a acetylation, suggesting inhibition of sirtuin1 and impaired calcium handling, which likely underlies the observed effects of high-dose NAM on arrhythmia and cardiac dysfunction.

**Conclusions:** NAM exhibits a narrow therapeutic window in aging-related cardiac dysfunction and arrhythmia, with its efficacy highly dependent on dose, duration, and biological context. While a moderate dose in late-life may be protective, chronic or excessive intake of NAM can induce arrythmia and impair cardiac function, likely through disruption of the SERCA2a activity. These findings underscore the importance of cautious and context-specific application of NAM in clinical settings.

## Introduction

Aging, a major risk factor for cardiac diseases, drives the progressive decline of cardiac function.^1^ With the increase in elderly in populations, the prevalence of aging-associated cardiac diseases, such as atrial fibrillation (AF)^2^ and heart failure^3^, has increased dramatically, leading to a substantial economic burden on healthcare systems. Aging is also associated with a reduction in nicotinamide adenine dinucleotide (NAD⁺) levels,^4^ a critical coenzyme involved in energy metabolism, redox balance, and signaling pathways essential for cardiac function.^5,6^ Lower levels of NAD^+^ have been implicated in increased AF susceptibility and cardiac dysfunction.^7,8^ NAD^+^-boosting compounds thus represent attractive therapeutic candidates for aging-related arrhythmogenesis and cardiac dysfunction.

Nicotinamide (NAM) is a form of vitamin B3 and a key precursor for NAD^+^ biosynthesis.^9^ Supplementation with NAM has shown protective effects in aging-related cardiometabolic disorders, including AF and heart failure. A previous study revealed that NAM ameliorated aging-induced diastolic dysfunction in 24-month-old mice by normalizing NAD^+^ levels.^10^ Our previous data showed that 10 mM NAM reduces AF-induced contractile dysfunction in tachypaced HL-1 cardiomyocytes and *Drosophila* models,^11^ though it remains unclear whether similar beneficial effects could be recapitulated in aging-related AF in mammalian models. In addition, emerging evidence suggests that the timing and duration of NAM intervention also influence its therapeutic outcomes. For instance, post-ischemic NAM intervention improved ischemic stroke, while longer pre-ischemic exposure worsened infarct size and induced neurological deficits.^12^ Similarly, in a rat brain injury model, NAM improved cognitive recovery only when delivered within 4 hours post-injury, with little benefit beyond that time window.^13^ These findings underscore the importance for a more comprehensive understanding of optimal dosing regimens to maximize the therapeutic potential of NAM.

While low to moderate doses are often beneficial, growing concerns have been raised regarding the possible adverse effects of high-dose NAM and the risks of long-term use.^14^ NAM concentrations above 20 mM have been reported to induce cytotoxicity and cell death.^15^ Dietary intervention with up to 32.77 mM NAM/kg caused detrimental metabolic and epigenetic changes in developing rats.^16^ These adverse effects of NAM may be attributed, at least in part, to its inhibition on sirtuins (SIRT) deacetylases,^17^ which regulate function and stability of numerous enzyme targets.^18^ Recent studies have shown that SIRT1 inhibition increases the acetylation of sarcoplasmic/endoplasmic reticulum Ca²⁺-ATPase 2a (SERCA2a), a critical Ca²⁺ pump responsible for proper cardiac relaxation and contractility, which in turn impairs its activity and contributes to both systolic and diastolic dysfunction as well as arrhythmogenesis.^19,20^ Therefore, it is of particular interest to investigate whether the cardiac-specific adverse effects of high-dose NAM impact SERCA2a activity by acetylation.

To address the above mentioned research gaps, we here study the effect of previously established 10 mM NAM on aging-associated cardiac arrhythmia in mouse hearts as a mammalian model, and further explore the potential cardiac-specific (adverse) effects of higher NAM doses using *ex vivo* mouse hearts, cardiomyocyte, and adult *Drosophila* models. Our findings reveal that long-term NAM intervention or concentrations exceeding 20 mM increase arrhythmia incidence and exert detrimental effects on cardiac function, at least partially by impairing SERCA2a activity through enhanced acetylation. This raises concerns regarding the safe and effective applications of NAM, particularly in aging contexts, and highlights the need for careful dose and duration considerations in both experimental research and potential clinical use.

## Methods

### HL-1 cardiomyocyte culture and contractile function

HL-1 cardiomyocytes were cultured in cell culture flasks or 6-well plates coated with 0.02% gelatin (G1393, Sigma) in a humidified atmosphere containing 5% CO₂ at 37 °C. The cells were maintained in complete Claycomb medium (51800C, Sigma) supplemented with 10% fetal bovine serum (TMS-016-B, Sigma), 100 U/mL penicillin-streptomycin (15140122, Thermo Fisher), 2 mM L-glutamine (25030024, Thermo Fisher), and 100 µM norepinephrine (A0937, Sigma).

The arrhythmia index, defined as the standard deviation of the cycle length normalized to the median cycle length, was determined by analyzing the spontaneous contractions of HL-1 cardiomyocytes, without electrical stimulation. Contractility parameters including cell shortening (peak amplitude change), time to peak, and peak to baseline duration of HL-1 cardiomyocytes were measured under electrical field stimulation (40 V, 1 Hz, 10 ms pulse duration) using a C-Pace100 culture pacer (IonOptix). In both cases, high-speed (120 frames per second) optical recordings of beating cells were acquired and analyzed using IonWizard 7.8 software (IonOptix). Data analysis was performed using CytoSolver software (IonOptix).

Prior to contractility measurements, HL-1 cardiomyocytes were treated with NAM (N0636, Sigma) at concentrations of 0, 10, 20, 50, or 100 mM for 4 h. Since a 1 h incubation with 100 mM NAM abolished spontaneous contractions (Video S1), only concentrations of 0, 10, 20, or 50 mM NAM were included for the 4 h for arrhythmia index assessment.

### Drosophila lifespan

Wild-type W1118 flies were incubated at 25 °C on a 12-h light/dark cycle. A rearing food containing (in g/L) 17 agar (76050048.5000, Boom), 54 glucose (G8270, Sigma), 26 yeast (903312, MP Biomedicals) and 1.3 nipagin (H3647, Sigma) was used as control food.

Within 8 h after eclosion, virgin female and male flies were separated and randomly distributed into a control or dietary intervention groups containing NAM at concentrations of 10, 20, 50, and 100 mM. Each group consisted of approximately 60-80 male and female flies, with 20 flies per tube. All flies were transferred to fresh food every 2 days, and the mortality was recorded.

### Dietary intervention and cardiac function measurement in *Drosophila*

To test the chronic effects of 10 mM NAM on arrhythmogenesis and cardiac function, the flies were fed either control or 10 mM NAM-supplemented diets at different times and durations, either starting from day 0 for 5 weeks (life-long) or starting at week 4 for 1 week (late-life). To test the detrimental effects of high-dose NAM on cardiac function, the flies were fed either control diet or 100 mM NAM-supplemented diet from day 0 for 1 week.

The cardiac function was measured in semi-intact *Drosophila* heart preparations using a previously established method.^21^ Briefly, after anesthetizing the flies with FlyNap (6353-173010, VOS instrument) for about 2 min, the heart tubes were exposed by removing the head and internal organs with microsurgery forceps. Dissections were performed in oxygenated artificial hemolymph containing 108 mM NaCl, 5 mM KCl, 2 mM CaCl_2_, 8 mM MgCl_2_, 15 mM HEPES, 1 mM NaH_2_PO_4_, 4 mM NaHCO_3_, 10 mM sucrose, and 5 mM trehalose, pH 7.1, at room temperature (RT). Heart activity was recorded for 20 s at 160 frames per second using a high-speed digital camera (DMK 33UX174, Imaging Source) mounted on a Leica DM IL LED microscope with a 10× lens and captured by IC Capture 2.5 software (The Imaging Source).

The cardiac physiology of the flies was assessed using a semiautomated optical heartbeat analysis program (SOHA) that quantifies functional parameters, including heart rate, diastolic and systolic intervals, contraction and relaxation phase intervals, fractional shortening, and shortening velocity, etc.^21,22^ Additionally, the arrhythmia index was calculated from the standard deviation of the heart period normalized to the median heart period. Incidence and burden of fibrillation events were evaluated based on M-modes cardiography records of heart edge movement over time, produced through SOHA analysis.

### *Ex vivo* mouse heart electrophysiological measurement

All the animal procedures were ethically approved by the Animal Welfare Committee of Wageningen University (2020.W-0019.009). B6JRccHsd(B6J)-*Nnt*^+^/Wuhap (*Nnt*^wt^) male and female mice, an in-house bred mouse line, were used for *ex vivo* electrophysiological measurement. Mice were heparinized and then sacrificed by cervical dislocation. Hearts were explanted, cannulated and connected to the Langendorff system (130102EZ-220, ADInstruments) perfused with modified Tyrode’s solution containing 130 mM NaCl, 4 mM KCl, 1.8 mM CaCl_2_, 1 mM MgCl_2_, 5.5 mM glucose, 1.2 mM NaH_2_PO_4_, and 24 mM NaHCO_3_, pH 7.4, at constant oxygenation with carbogen (SOL group). A water thermo-regulator was used to maintain constant temperature (37 ± 0.5 °C) for the complete Langendorff system. Cardiac electrocardiograms (ECG) were recorded using three electrodes (MLA1213 and MLA1212, ADInstruments). The positive electrode was placed beneath the apex of the left ventricle, while the negative and ground electrodes were positioned on the cannula (Figure 1A). Recordings were acquired with a bio-amplifier recording system (FE231, ADInstruments). Programmed electrical stimulation was applied using a stimulus isolator (FE180, ADInstruments) via a spring-loaded platinum electrode (140157M-STISO, AD Instruments) placed on the surface of the right atrium (Figure 1A). All data were acquired using the 16-channel PowerLab system (PL3516/P, ADInstruments).

**Figure 1.**
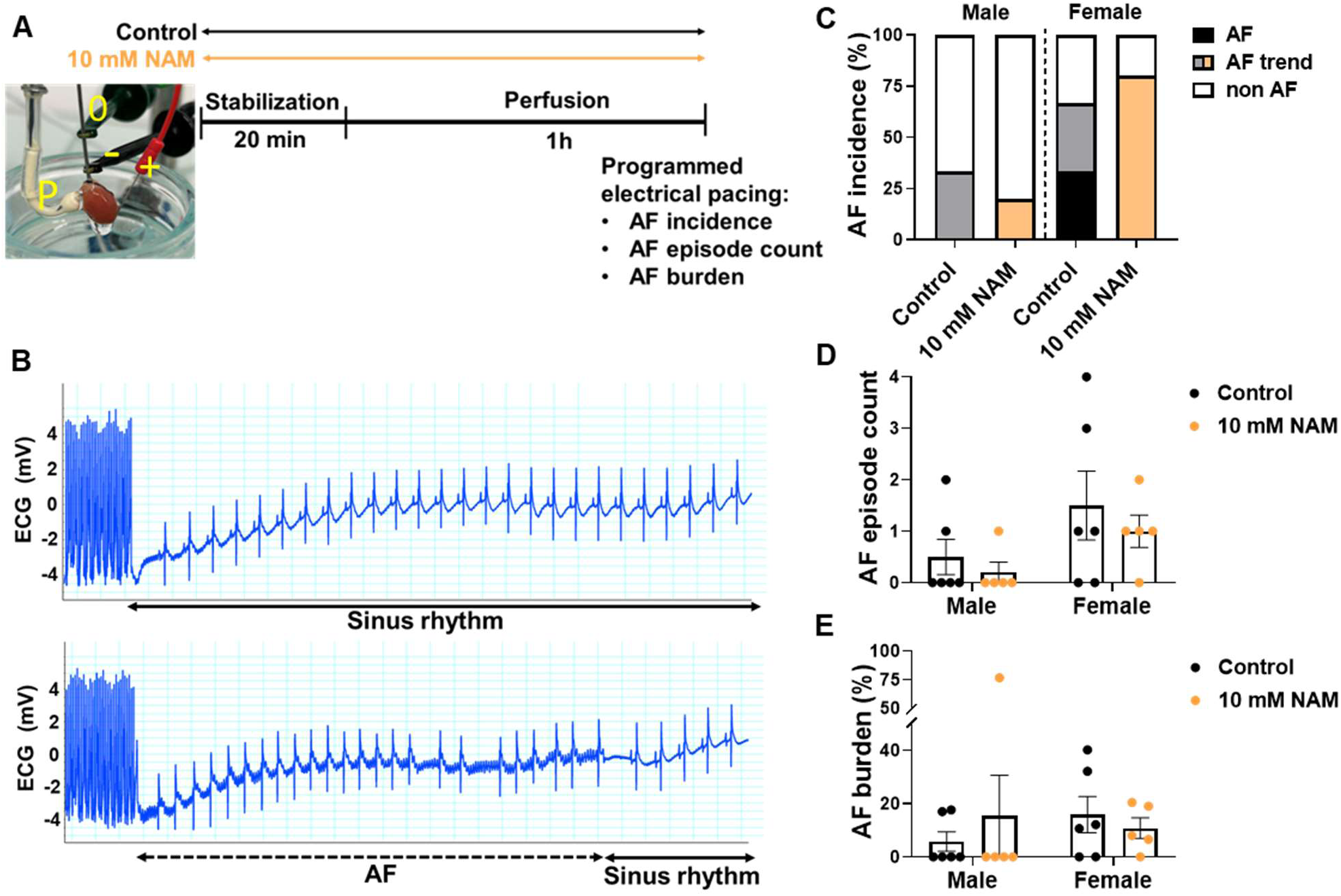
Acute perfusion with 10 mM nicotinamide (NAM) has no effect on aging-related atrial fibrillation (AF) in 18-month-old *ex vivo* mouse hearts. (A) Experimental scheme of Langendorff *ex vivo* mouse hearts perfused with control or 10 mM NAM-supplemented buffer. Electrocardiogram (ECG) was recorded with the positive electrode (‘+’) positioned beneath the apex of the left ventricle, and the negative (‘-’) and ground (‘0’) electrodes on the cannula. Programmed electrical stimulation was applied via a pacing electrode (‘P’) placed on the surface of the right atrium. (B) Representative ECG traces from AF-negative (top) and AF-positive (bottom) mouse hearts. (C-E) AF incidence, episode count and burden in hearts from the control (n=6 for both sexes) and 10 mM NAM (n=5 for both sexes). Data are expressed as mean ± standard error of the mean. Chi-square test was used for AF incidence and Scheirer-Ray-Hare test followed by Dunn’s post-hoc test was used for AF episode count and burden.

Hearts from 18-month-old (18M) male and female mice were used to test the effects of perfusion of 10 mM NAM. 12 hearts were perfused with the control modified Tyrode’s solution and 10 hearts were perfused with 10 mM NAM supplemented solution throughout the whole procedure. Each heart in the Langendorff system was stabilized for 20 min followed by a 1 h perfusion before the electrophysiological measurement. In the electrophysiological measurement, the pacing capture threshold was determined by applying series of 2 ms current pulses at 6 Hz and 8 Hz for consistency of stimulus capture. Then the AF susceptibility was assessed using programmed electrical stimulation as pacing is necessary to induce arrhythmia in isolated mouse hearts where spontaneous events are rare.^23,24^ The programmed electrical stimulation was performed using 10 s electrical burst pacing with cycle length of 20 ms (50 Hz) at a three-time capture threshold current for three times, with a 30 s recovery period between each assessment.^25^ AF was characterized by rapid and irregular atrial electrograms longer than 0.5 s.^26^ An AF positive heart is defined when AF occurs at least two out of three repeats, and an AF trend heart is defined when AF occurs one out of three repeats. AF burden was calculated as the cumulative AF duration divided by the total monitoring time.

Hearts from 6-month-old (6M) male and female mice were used to assess the effects of perfusion of 100 mM NAM (n=6), 100 mM glucose (n=5), or 50 mM NaCl (n=5), with the latter two serving as isosmotic controls. Each heart in the Langendorff system was stabilized for 20 min before switching to the respective perfusates until the onset of ventricular fibrillation (VF) or cardiac arrest. Heart tissue was then snap frozen and stored at −80 °C for further molecular analysis.

### NAD assay

Total NAD (NAD_t_: NAD^+^ and NADH) and NADH levels were measured according to the manufacturer’s instructions of the assay kit (ab65348, Abcam). In short, 6M and 18M old female mouse right atrial tissue were lysed in NADH/NAD Extraction Buffer (ab65348, Abcam) and centrifuged for 5 min at 4°C at 21300 × *g*. To measure NAD_t_, after equalizing the protein concentration of the supernatant, 50 µl of each sample was mixed with 100 µl reaction mix containing 98 µl NAD cycling buffer (ab65348, Abcam) and 2 µl NAD cycling enzyme mix (ab65348, Abcam), and incubated at RT for 5 min to convert NAD^+^ to NADH. 10 µl NADH developer buffer (ab65348, Abcam) was then added and incubated at RT for 3 h. To measure NADH, NAD^+^ in each sample was decomposed by incubation at 60 °C for 30 min before measurement. Absorbance was measured at 450 nm using a plate reader (235197, BioSPX) to determine NAD_t_ and NADH levels.

### Immunoprecipitation and Western blotting

Left ventricular tissue (n=3) from mouse hearts perfused with control perfusate or 100 Mm NAM was homogenized in immunoprecipitation (IP) buffer containing 50 mM Tris-HCl pH 7.4, 0.5% Triton X-100, 150 mM NaCl, 1 mM ethylenediaminetetraacetic acid, 1 μM trichostatin A, 10 mM NAM, and protease (04693159001, Roche) and phosphatase inhibitor cocktail (04906837001, Roche). The homogenates were centrifuged at 1000 × *g* for 10 min at 4°C. Equal amounts of protein (600 ug) were incubated with antibodies against Pan Acetylation (66289-1-Ig, Proteintech) or mouse IgG (I-2000-1, Vector laboratories) as negative control at 4 °C on a rotation wheel at 12 revolutions per minute (rpm) overnight. The homogenates were then incubated with protein-G/A agarose beads (88802, Fisher scientific) at 4 °C for 2 h. The resulting protein-beads complexes were washed three times with IP buffer before elution at 95 °C for 5 min with the elution buffer containing IP buffer mixed with 50 mM DTT and 1x NuPAGE LDS sample buffer (NP0007). For Western blotting analysis, proteins were separated by SDS-PAGE (NP0323BOX, Invitrogen) at 110 V for 30 min and then 150 V for 60 min in XCell SureLock system (EI0001, Novex) and transferred to a methanol-activated PVDF membrane (IPFL85R, Sigma) at 300 mA on ice for 1.5h. The membrane was then blocked with Intercept Blocking Buffer (927-60001, LI-COR) for 1 h at RT, and incubated with primary antibodies against SERCA2a (1:1000 dilution, 4388, cell signaling), and Pan Acetylation (1:2000 dilution, 66289-1-Ig, Proteintech) overnight at 4°C. The membrane was subsequently incubated with secondary antibodies IRDye 680RD Donkey anti-rabbit (1:5000 dilution, 926-68073, LI-COR) or IRDye 800CW Donkey anti-mouse (1:5000 dilution, 926-32212, LI-COR) for 1 h at RT, and visualized using an Odyssey scanner (LI-COR).

### Statistical analysis

All the comparisons and graphs were made using the GraphPad Prism software (v9.5.1). AF incidence of *ex vivo* mouse hearts or fibrillation event incidence of *Drosophila* were tested using the Chi-square test. AF episode count and burden of *ex vivo* mouse hearts or fibrillation burden of *Drosophila* were tested using the Scheirer-Ray-Hare test followed by Dunn’s post-hoc test. The effects of 10 mM or 100 mM NAM on cardiac function in *Drosophila* were tested by two-way analysis of variance (ANOVA) followed by post-hoc Fisher’s LSD test. Contractility among different doses of NAM in HL-1 cardiomyocytes was tested by one-way ANOVA followed by post-hoc Fisher’s LSD test. Differences between survival curves of *Drosophila* were analyzed by the log-rank test. The effects of high-dose NAM on SERCA2a acetylation in *ex vivo* mouse hearts and NAD levels were tested by two tailed unpaired Student t-test. The statistical significance was shown as \**P* < 0.05, \*\**P* < 0.01, \*\*\**P* < 0.001, \*\*\*\**P* < 0.0001.

## Results

### Acute treatment of 10 mM NAM did not prevent aging-related AF in *ex vivo* 18M-old mouse hearts

Our previous study showed that 10 mM NAM protected against tachypacing-induced contractile dysfunction in HL-1 cardiomyocytes and *Drosophila*, primarily by restoring NAD^+^ levels.^7,11^ Since aging is the most common risk factor for AF and is known to reduce NAD^+^ levels,^27,28^ we further studied whether 10 mM NAM could also protect against aging-related AF using isolated 18M-old mouse hearts on the Langendorff system with programmed electrical pacing (Figure 1A and 1B). We found that hearts from 18M-old mice exhibited a decline of NAD^+^ and NAD^+^/NADH ratio (Figure S1), and displayed a considerable incidence of pacing-induced AF, especially in females (∼33.3 % AF positive and ∼33.3 % AF trend, Figure 1C). However, supplementation of 10 mM NAM in the perfusate did not result in significant changes in AF incidence, episode count or burden compared to controls (Figure 1B-E). Unlike previous findings with longer durations of NAM treatment in different model systems,^7,11^ 1-h acute perfusion of 10 mM NAM did not alleviate AF in *ex vivo* hearts from aged mice.

### Effects of different timing and intervention durations of 10 mM NAM on aging-related arrhythmia and contractile dysfunction in *Drosophila*

Since acute perfusion of 10 mM NAM failed to attenuate pacing-induced AF in old *ex vivo* mouse hearts, we next investigated whether different timing and intervention durations could influence aging-related arrhythmia in an *in vivo* setting. Employing *Drosophila melanogaster* as a recognized model for aging,^29^ we treated flies with 10 mM NAM diet either for 1 week prior to measurements or for 5 weeks throughout their lifespan before assessment (Figure 2A). Our results showed that 5-week-old fly hearts recapitulated aging-induced arrhythmia and cardiac contractile dysfunction, characterized by prolonged diastolic and systolic intervals, relaxation and contraction phase intervals, and increased arrhythmia index, as well as reduced heart rate and shortening velocity compared to 1-week-old control flies (Figure 2B-H, S2A, S2B). Interestingly, short-term (1-week in week 5) dietary intervention of 10 mM NAM ameliorated aging-induced prolongation of diastolic intervals (Figure 2C) and the arrhythmia index (Figure 2E), and showed a nonsignificant trend toward rescuing the reduced shortening velocity (*P* = 0.15, Figure 2H), exclusively in male flies, without significant influences on systolic intervals (Figure 2D) or heart rate (Figure 2F). Surprisingly, a life-long (5 weeks) intervention with 10 mM NAM did not alleviate aging-induced arrhythmia in male flies and even increased the arrhythmia index in female flies (Figure 2E), despite an increase in heart rate in both sexes compared to 5-week-old flies on the control diet (Figure 2F). Additionally, the fibrillation burden following 5 weeks of 10 mM NAM treatment was greater than that observed with the 1-week intervention in week 5 in female (Figure S2F), but not in male flies (Figure S2E). No significant differences in fractional shortening (Figure 2G) and fibrillation event incidence (Figure S2C and S2D) were found among all groups. Together, these findings suggest that shorter-term in late-life, but not life-long, intervention with 10 mM NAM alleviates aging-induced arrhythmia and contractile dysfunction in *Drosophila* in a sex-specific manner.

**Figure 2.**
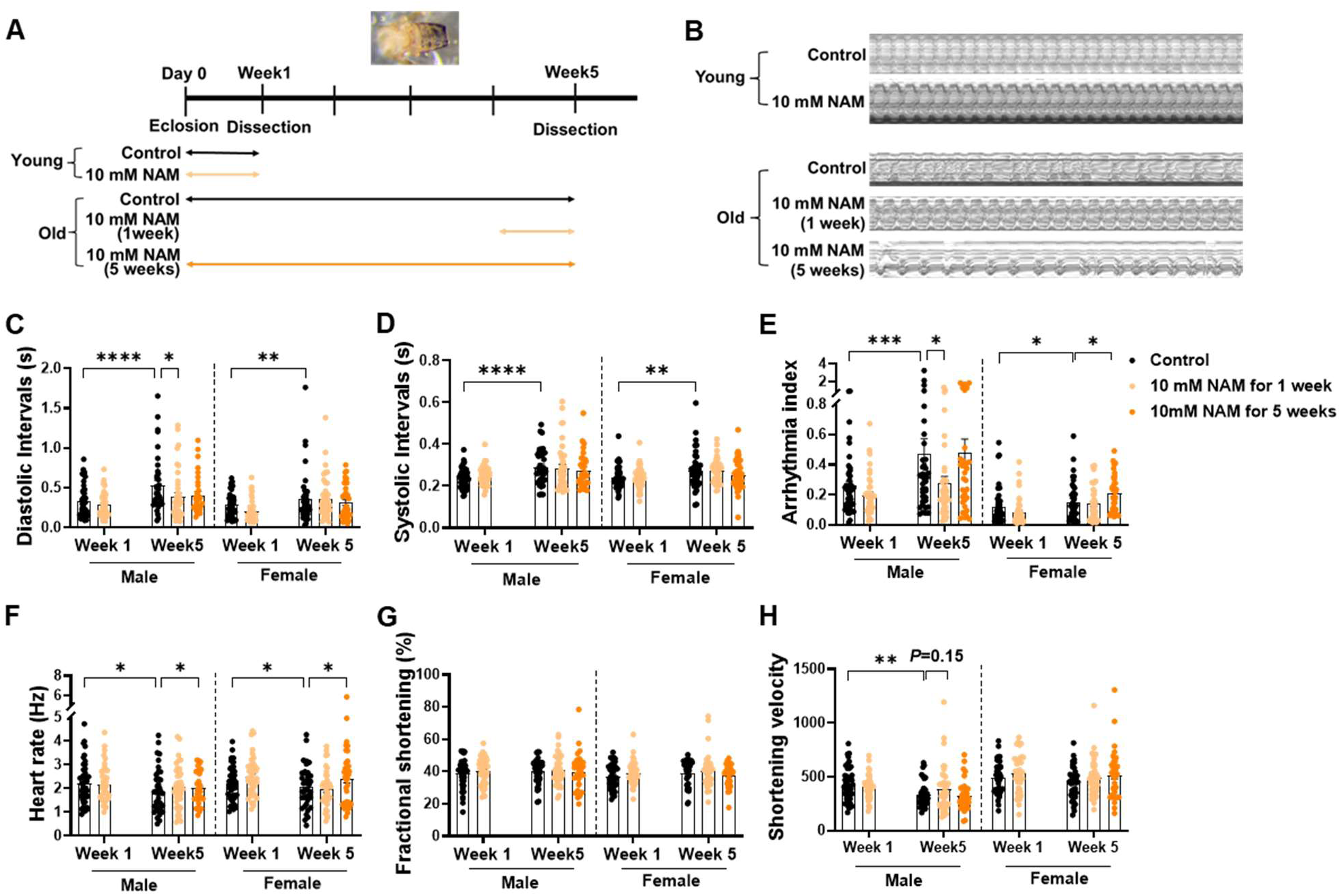
Shorter-term in late-life, but not life-long intervention with 10 mM nicotinamide (NAM) alleviates aging-induced arrhythmia and contractile dysfunction in *Drosophila melanogaster*. (A) Experimental design with control or 10 mM NAM-supplemented diet for one week (in week 1 and week 5) or five weeks. (B) Representative 20-s M-mode cardiograms of semi-intact *Drosophila* heart preparations. (C) Diastolic intervals, (D) systolic intervals, (E) arrhythmia index, (F) heart rate, (G) fractional shortening, and (H) shortening velocity of young flies fed with a control diet (n=45 males, 41 females) or a 10 mM NAM-supplemented diet (n=45 males, 44 females), and of old flies fed with a control diet (n=38 males, 39 females), a 10 mM NAM-supplemented diet for 1 week (n=40 males, 39 females), or for 5 weeks (n=33 males, 37 females). Arrythmia index=SD/Median heart period. Data are expressed as mean ± standard error of the mean. Two-way ANOVA followed by post-hoc Fisher’s LSD test was used. \**P* < 0.05, \*\**P* < 0.01, \*\*\**P* < 0.001, \*\*\*\**P* < 0.0001.

### A dose-dependent effect of NAM on arrhythmia incidence and contractile function of HL-1 cardiomyocytes

Given that a long-term intervention with 10 mM NAM exhibited pro-arrhythmic effects in female flies (Figure 2E), we hypothesized that higher doses of NAM might similarly exert adverse effects on the arrhythmia incidence and cardiac contractile function. To test this, we determined the impact of a series of high NAM doses on arrhythmia index and cardiac contractility in HL-1 cardiomyocytes. Indeed, NAM concentration above 20 mM led to a gradual increase in arrhythmia index of spontaneous beating HL-1 cardiomyocytes (Figure 3A and 3B). Notably, 100 mM NAM treatment totally abolished the spontaneous contractions of HL-1 cardiomyocytes (Video S1). In addition, NAM exhibited a dose-dependent effect on cellular contractility under pacing where 10 mM NAM had no significant effects compared to the control, whereas higher doses (20, 50 and 100 mM) progressively reduced contractility as measured by cell shortening percentage (Figure 3C and 3D), and shortened systolic phase and diastolic phase contraction duration (Figure 3E and 3F). Collectively, these results demonstrate a dose-dependent effect of NAM on HL-1 cardiomyocytes, with higher doses causing arrhythmia and exerting detrimental effects on contractility.

**Figure 3.**
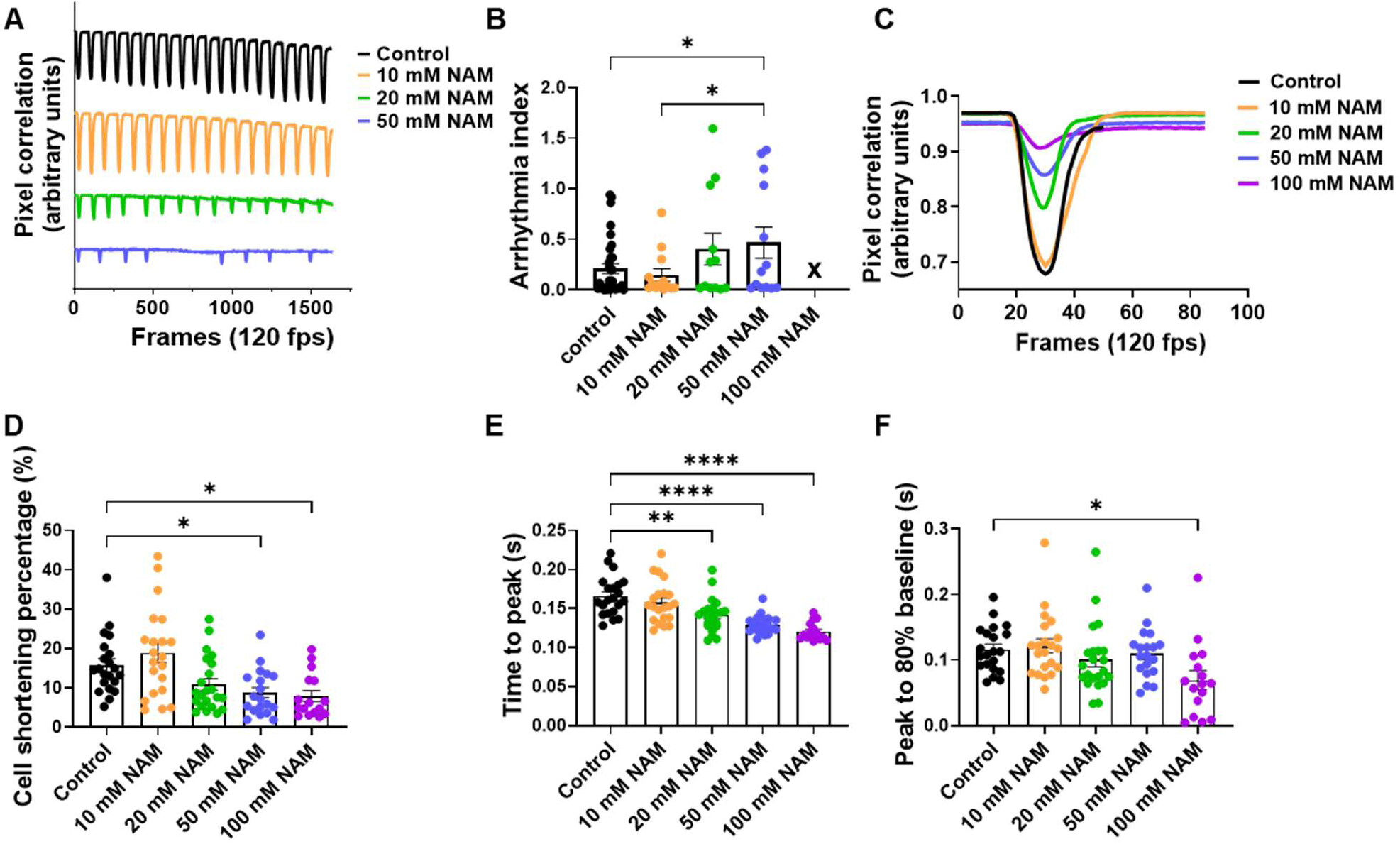
A dose-dependent effect of nicotinamide (NAM) on arrhythmia and contractile function of HL-1 cardiomyocytes. (A) Representative contractile traces of HL-1 cardiomyocytes treated with different doses of NAM without pacing recorded at 120 frames per second (fps). (B) Quantified arrhythmia index of spontaneous beating cells incubated with 0 mM (control; n=33), 10 mM (n=12), 20 mM (n=12), or 50 mM (n=13) of NAM for 4 h. Arrythmia index=SD/Median heart period. 100 mM NAM is not shown due to abolished spontaneously contraction of the HL-1 cardiomyocytes. (C) Representative contractile traces of HL-1 cardiomyocytes assessed under 1 Hz pacing (40 V, 10 ms pulse duration) incubated with 0 mM (control), 10 mM, 20 mM, 50 mM and 100 mM for 4 h. Quantification of (D) cell shortening percentage, (E) time to peak, (F) time of peak to 80% baseline for the control (n=21), 10 mM (n=21), 20 mM (n=22), 50 mM (n=19), and 100 mM (n=16) NAM-treated cells. Data are expressed as mean ± standard error of the mean. One-way ANOVA followed by post-hoc Fisher’s LSD test was used. \**P* < 0.05, \*\**P* < 0.01, \*\*\*\**P* < 0.0001.

### A dose-dependent effect of NAM on the lifespan of *Drosophila*

To evaluate whether the dose-dependent effects of NAM on cardiac function observed in HL-1 cardiomyocytes could be translated to a whole-body context, we assessed the lifespan of *Drosophila* fed the same range of NAM doses. In line with prior *in vitro* results (Figure 3), 10 mM NAM intervention prolonged the lifespan of female flies but not males (Figure 4, Table S1). In females, the hazard ratio was 0.6039 (95% confidence interval 0.441 to 0.827), which indicated a decreased mortality risk relative to the control group (Table S1). However, higher concentrations, i.e. 20, 50, and 100 mM, significantly reduced lifespan in both sexes (Figure 4), with the hazard ratios all significantly higher than 1, indicating an increased mortality risk compared with the control group (Table S1). Notably, the sensitivity to higher doses of NAM appeared to be sex-specific. At 20 mM, both male and female flies showed reduced life span, with males showing a more rapid mortality after ∼50 days compared to females (Figure 4). Consistently, 50 mM NAM caused a shortened lifespan of both sexes, with an early and rapid loss of male flies, while female survival was less affected initially (Figure 4). Most strikingly, exposure to 100 mM NAM led to death in both sexes within approximately 1 week (Figure 4). Overall, these findings demonstrate a dose-dependent effect of NAM on *Drosophila* lifespan, with males more susceptible to adverse effects of doses above 10 mM.

**Figure 4.**
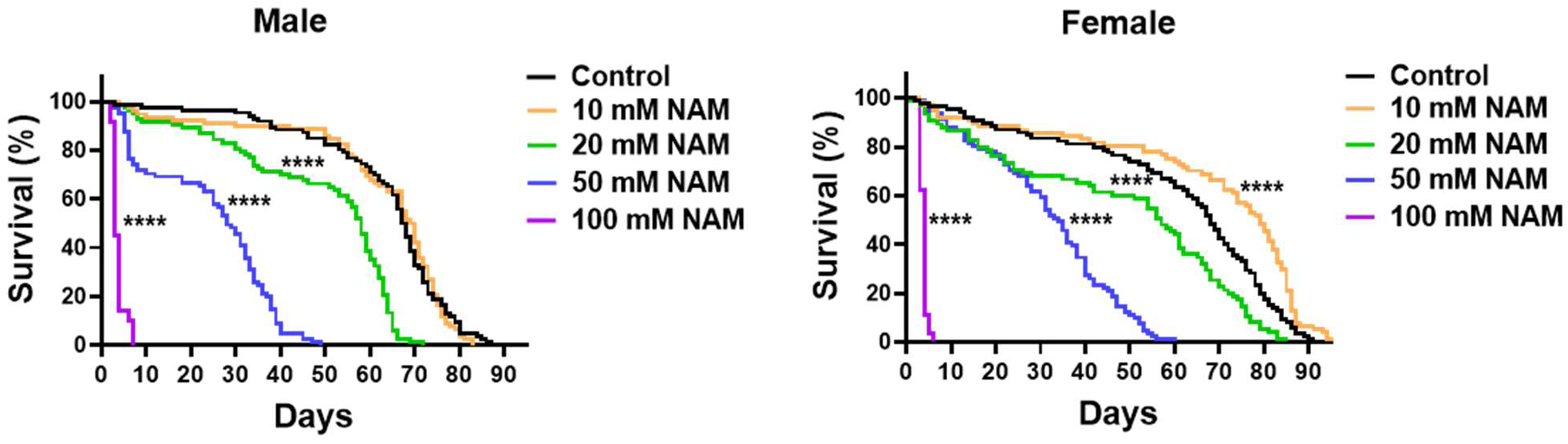
A dose-dependent effect of nicotinamide (NAM) on lifespan of *Drosophila melanogaster*. Survival curves of male and female flies were fed with diet supplemented with 0 mM (control; n=86 males, 85 females), 10 mM (n=78 males, 77 females), 20 mM (n=72 males, 75 females), 50 mM (n=81 males, 81 females), and 100 mM (n=71 males, 80 females) NAM. Log-rank test was used to analyze the differences between survival curves of flies. \*\*\*\**P* < 0.0001.

### High-dose NAM reduced heart rate and induced arrhythmia in *Drosophila*

Since higher doses of NAM, particular 100 mM, led to near-complete mortality in *Drosophila* within 1 week, we further checked its cardiac-specific effects in the few surviving flies at the end of the 1-week exposure. Remarkably, 100 mM NAM severely impaired cardiac rhythm (Figure 5A), as indicated by an increased diastolic intervals (Figure 5B) and decreased heart rate (Figure 5D), with systolic intervals unaffected (Figure 5C) in both sexes. Arrhythmia index was increased in both male and female flies (Figure 5E), while the incidence and burden of fibrillation events were elevated significantly in males, but not in females (Figure 5F-5H). However, the contraction and relaxation phase intervals, fractional shortening, and shortening velocity were not significantly influenced by 1-week intervention with 100 mM NAM (Figure S3A-S3D). Taken together, these findings demonstrate that 100 mM high-dose NAM reduces physiological heat rate, induces cardiac arrythmia and fibrillation in *Drosophila,* which might underlie its detrimental effect on the *Drosophila* lifespan (Figure 4).

**Figure 5.**
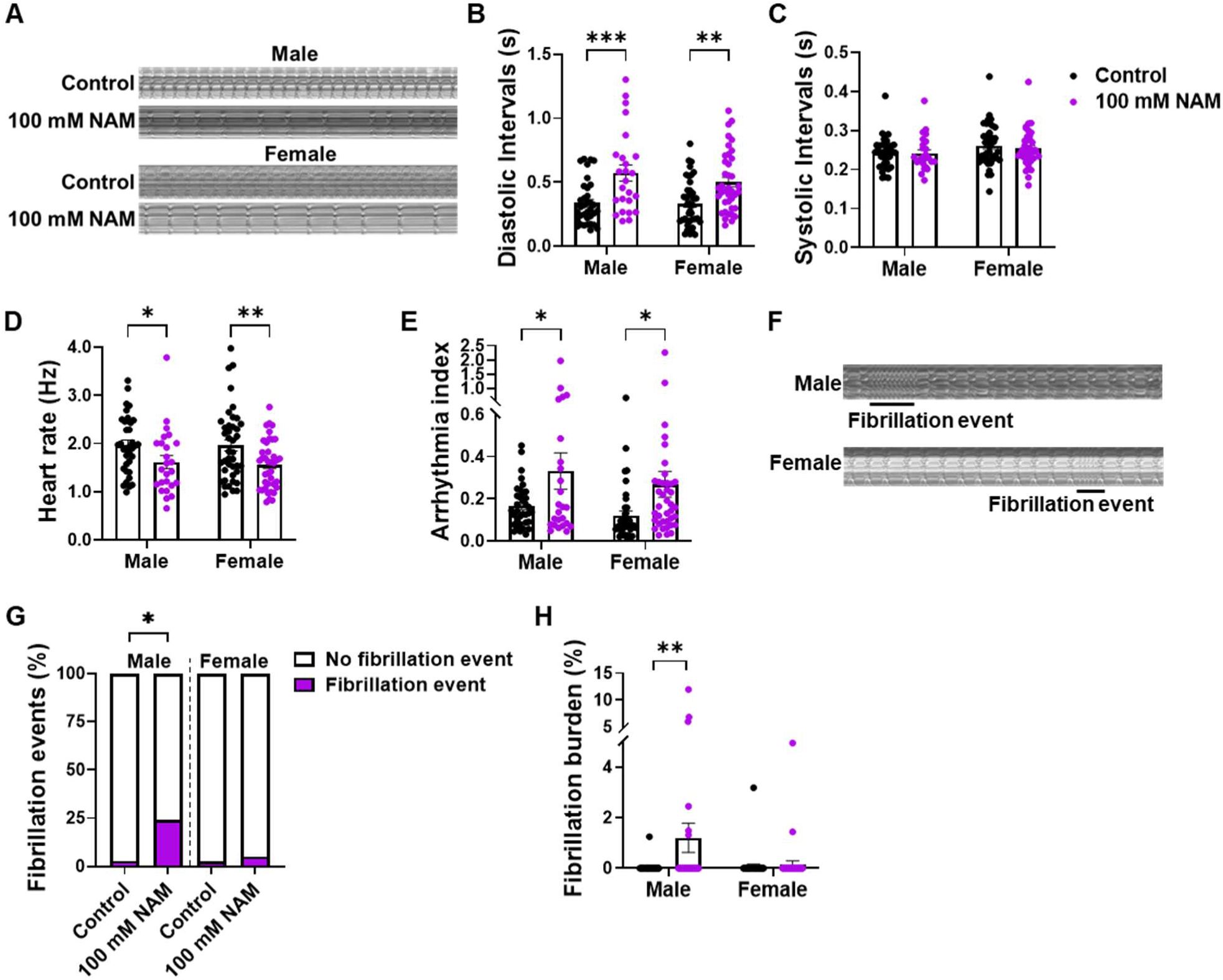
High-dose nicotinamide (NAM) leads to cardiac arrhythmia in *Drosophila melanogaster*. (A) Representative 20-s M-mode cardiograms of semi-intact heart preparations of flies fed with diet supplemented with 0 mM NAM (control; n=35 males, 41 females) or 100 mM NAM (n=25 males, 39 females). Quantification of (B) diastolic intervals, (C) systolic intervals, (D) heart rate (E) arrhythmia index. Arrythmia index=SD/Median heart period. Two-way ANOVA followed by post-hoc Fisher’s LSD test was used. (F) Representative 20-s M-mode cardiograms, and quantification of (H) fibrillation event incidence and (I) fibrillation burden in the control and 100 mM NAM groups. Chi-square test was used for fibrillation events and Scheirer-Ray-Hare test followed by Dunn’s post-hoc test was used for fibrillation burden. Data are expressed as mean ± standard error of the mean. \**P* < 0.05, \*\**P* < 0.01, \*\*\**P* < 0.001.

### High-dose NAM induced ventricular fibrillation in *ex vivo* mouse hearts partly through regulating SERCA2a acetylation

We finally investigated the detrimental effects of high-dose NAM on arrhythmia incidence and cardiac function in a mammalian model for further insight. After 20 min of stabilization, 100 mM NAM was perfused into the *ex vivo* Langendorff mouse heart, and ECG was monitored (Figure 6A). Strikingly, the acute perfusion with 100 mM NAM rapidly triggered VF events, impaired contractile function, and eventually resulted in cardiac arrest in all 6 tested hearts (Figure 6B, S4B). In contrast, perfusion with an equivalent osmotic molar concentration of glucose or NaCl did not cause any VF in any of the 5 hearts tested (Figure S4), suggesting that the VF induction was specific to NAM. Since NAM is a known inhibitor of SIRT deacetylases, we hypothesized that high-dose NAM might impair the deacetylation activity of SIRT, thereby influencing its target protein SERCA2a, whose dysfunction is closely associated with cardiac arrhythmia through its central role in calcium handling and electrical stability.^19,20^ Indeed, hearts perfused with 100 mM NAM exhibited a trend toward increased levels of acetylated SERCA2a (*P* = 0.0774, Figure 6C, 6D), indicating a reduced SERCA2a-mediated calcium uptake activity.^19^ This poteinally explains, at least in part, the occurrence of VF events in 100 mM NAM-treated *ex vivo* mouse hearts.

**Figure 6.**
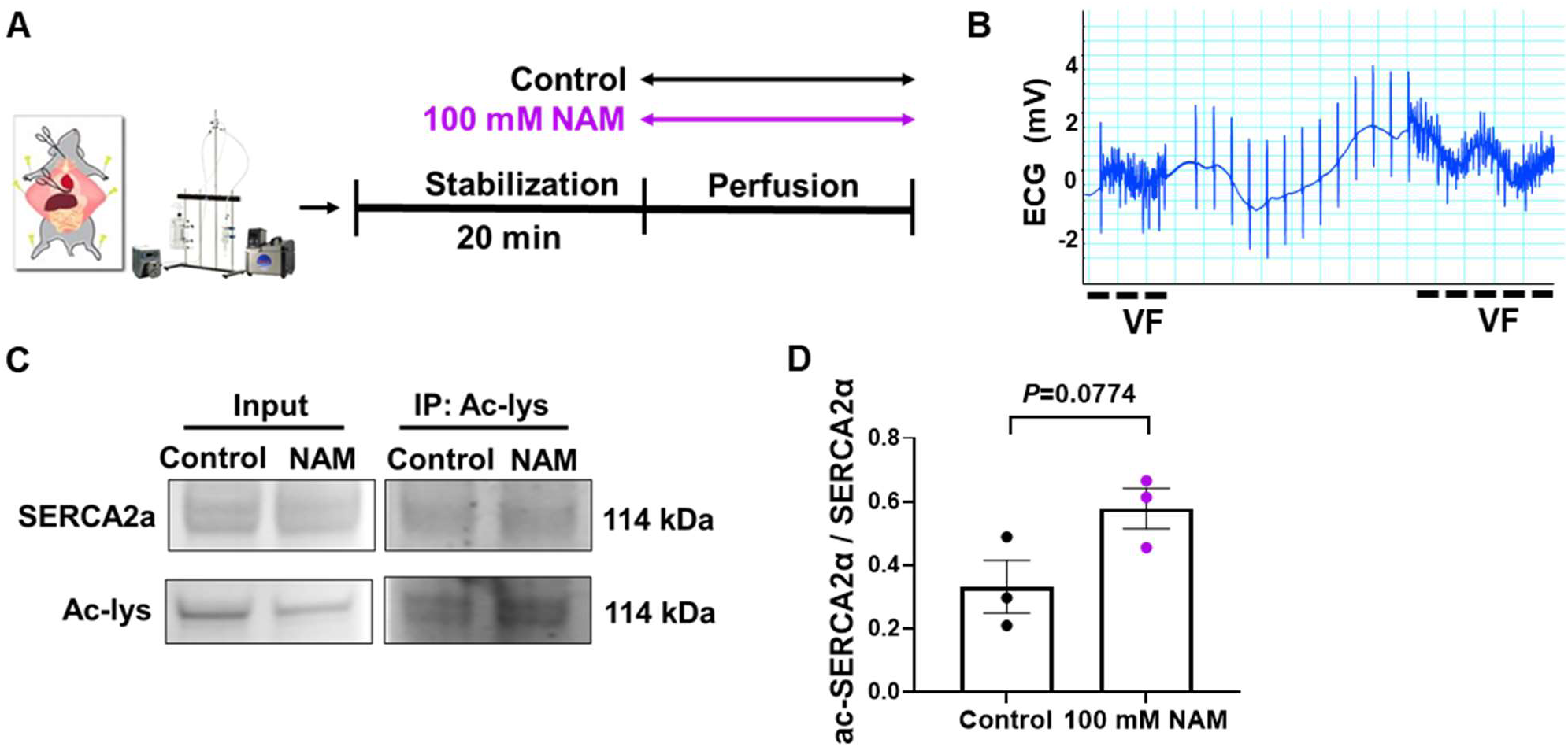
High-dose nicotinamide (NAM) causes ventricular fibrillation (VF) in *ex vivo* 6-month old mouse hearts partly through sarcoplasmic/endoplasmic reticulum Ca²⁺-ATPase 2a (SERCA2a) acetylation. (A) Experimental scheme of Langendorff *ex vivo* mouse hearts perfused with 0 mM NAM (control) or 100 mM NAM-supplemented buffer. (B) Representative electrocardiogram (ECG) shows that acute perfusion with 100 mM NAM rapidly induced VF, contractile dysfunction, and eventually led to cardiac arrest (n=4 males, 2 females). (C) Representiative blots and (D) quantification of acetylated SERCA2a levels of left venticular tissue from control and 100 mM NAM-perfused *ex vivo* mouse hearts (n=3). Data are expressed as mean ± standard error of the mean. Two tailed unpaired Student t-test was used.

## Discussion

In the current study, we investigated the effects of NAM on aging-related arrhythmia and cardiac dysfunction, and explored the cardiac-specific adverse effects of high NAM doses and a plausible contributing mechanism. Taking *ex vivo* mouse hearts and *in vivo Drosophila* models together, we found that 10 mM NAM showed at most modest cardioprotection. Short-term treatment during the late-life phase in male flies mitigated arrhythmia and contractile dysfunction, while long-term treatment prevented aging-induced reduction in heart rate but exhibited pro-arrhythmic effects. In contrast, higher concentrations of NAM (>10 mM) dose-dependently induced arrhythmia and contractile dysfunction in HL-1 cardiomyocytes, reduced lifespan, and promoted cardiac dysfunction and arrhythmia/fibrillation in *Drosophila*, with males displaying greater susceptibility to these toxicity effects. Consistently, high-dose NAM perfusion (100 mM) led to acute VF in *ex vivo* mouse hearts, potentially mediated by acetylation of SERCA2a. These findings suggest that the cardiac effects of NAM are highly dependent on dose, timing, duration, and modified by sex. Supplementation with NAM has been proposed to promote healthy aging in various models by elevating NAD^+^ levels. For instance, 0.2 to 1 mM NAM extended the lifespan of *C. elegans* in a dose-dependent manner,^30^ while 1 to 2 mM NAM protected against oxidative stress and improved mitochondrial function in both cellular and *Drosophila* models.^31^ However, in our study, acute perfusion with 10 mM NAM did not significantly alter AF susceptibility in aged mouse hearts. This was unexpected given our previous evidence supporting its anti-arrhythmic effects in tachypaced HL-1 cardiomyocytes and *Drosophila* models.^11^ One possible reason is likely that a 1-h acute NAM perfusion may not be sufficient to restore the declined NAD⁺ level in the aged mouse heart.^27,28^ The slower uptake of NAM and reduced enzymatic activity in aged tissue may delay NAD⁺ biosynthesis.^32^ Another explanation could be the complex pathophysiological basis of aging-related AF. Tachypacing mimics AF partially by inducing oxidative DNA damage, which acutely depletes NAD^+^ due to poly(ADP)-ribose polymerase 1 (PARP1) activation.^7^ However, aging-associated AF may involve more complex structural remodeling and chronic oxidative stress beyond NAD⁺ depletion,^33^ where a single acute intervention of NAM maybe insufficient to reverse NAD^+^ depletion and its chronic and multifactorial pathology.

Clinically, NAM at doses ranging from 1 to 5 g/day has been shown to be safe in humans. This dosage corresponds to an immediate post-dose plasma concentration of roughly 1.64 to 8.2 mM NAM, assuming a blood volume of approximately 5 L for a 70 kg adult. However, it is well recognized that the public use of nutritional supplements often exceeds recommended daily doses. Therefore, assessing the potential cardiac side effects of high-dose of NAM is clinically relevant. In this study, we tested supraphysiological NAM doses and found that higher doses of NAM displayed significant whole-body and cardiac-specific adverse effects. NAM doses above 20 mM increased arrhythmia index and impaired contractile function in HL-1 cardiomyocytes, and reduced lifespan in both male and female *Drosophila*. 100 mM NAM even abolished spontaneous contraction in HL-1 cardiomyocytes, reduced heart rate and caused cardiac fibrillation in *Drosophila*, as well as induced VF events in *ex vivo* mouse hearts. Similar adverse effects of NAM have been reported previously. Excess NAM was linked with neurodegeneration in Parkinson’s disease,^34^ and chronic high-dose NAM supplementation increased the risk of adverse outcomes in individuals predisposed to cerebral ischemia.^12^ High-level of NAM can exert negative effects through multiple pathways. Notably, NAM is a noncompetitive inhibitor of SIRT1 deacetylase, which regulates broad aspects of cellular signaling and homeostasis.^18^ Even low dose (50 µM) of NAM has been shown to sufficiently inhibit SIRT1 activity *in vitro.*^17^ Previous studies demonstrated that downregulation of SIRT1 increased acetylation of SERCA2a, which in turn led to reduced SERCA2a activity, impaired calcium handling, and cardiac defects in both HL-1 cardiomyocytes and mice.^19^ Consistent with these findings, we found increased SERCA2a acetylation in mouse hearts perfused with 100 mM NAM, reflecting impaired SERCA2a activity, which may explain the occurrence of VF and cardiac dysfunction. Regardless of the mechanisms, these data highlight previously unrecognized cardiac risks of chronic high-dose NAM, particularly for individuals predisposed to cardiovascular diseases such as AF and heart failure. Since most vitamin B3 supplements, NAM, nicotinic acid (NA), nicotinamide riboside (NR), and nicotinamide mononucleotide (NMN), circulate as NAM,^35^ this warrant caution in taking high-dose vitamin B3 supplementation and, especially, in applying intravenous NAD^+^ therapy.

Apart from dosage, the efficacy of NAM may also depend on the intervention timing and duration. We observed that only short-term intervention of 10 mM NAM, initiated at a late stage of life, protected against aging-induced cardiac arrhythmia and contractile dysfunction, but life-long intervention did not show protection in male flies and even elevated arrhythmias in females. A comparable timing-dependent adverse effect was reported in a study using rats, where 500 mg/kg NAM significantly ameliorated acetaminophen-induced biochemical changes and pathological injuries, but induced hepatotoxicity when given to healthy animals or prior to intoxication.^36^ These adverse effects of NAM seem to occur only when NAM is supplied at a young age or under healthy conditions, when NAD^+^ levels are normally unaffected. In such contexts, NAM supplementation may be excessive and accumulate chronically, leading to side effects independent of NAD⁺-mediated mechanisms. These results are in line with our previous results where high-dose NR intervention impaired metabolic flexibility, reduced insulin sensitivity and exacerbated inflammation in young healthy male mice fed on obesogenic diet.^37,38^ On the contrary, treatment of NAM at old age, when NAD^+^ levels are often reduced, could yield beneficial outcomes. These contrasting effects underscore the importance of therapeutic window for NAM intervention, especially in the context of aging and cardiovascular diseases.

Sex-specific responses were also observed in the sensitivity to NAM. Exclusively in male flies, 10 mM NAM alleviated aging-induced cardiac arrhythmia, while 20 mM and 50 mM led to more rapid mortality rate and 100 mM NAM induced fibrillation events, suggesting that males are more sensitive to the effects of NAM, either at moderate or extreme high doses. Such sexually dimorphic responses were also reported in a mouse model of Friedreich’s ataxia, where male mice exhibited more severe cardiac NAD⁺ depletion and greater improvement in cardiac phenotypes with NAD⁺ precursor treatment, whereas females maintained NAD⁺ levels and showed less pronounced benefit.^39^ Similarly, a previous study found NAM alleviated heart failure in aged male mice.^10^ Our previous studies further showed that male mice are more susceptible than females to both vitamin B3 deficiency^40^ and the adverse effects of high-dose NR supplementation.^37^ In this study, the protective effects of 10 mM during late-life in male flies seem to be due to a higher arrhythmia incidence during aging compared with females. However, previous animal and clinical studies have shown inconsistent sex differences in arrhythmia incidence. While some studies reported that males are generally more susceptible to AF than females,^41^ a study conducted in an aged population suggested that women are at higher risk for AF incidence than men for a given height and weight.^42^ Further investigation into sex difference in NAM efficacy for aging-related cardiac dysfunction and arrhythmia is therefore warranted.

## Conclusions

Our results revealed a dose- and context-dependent profile of NAM in cardiac aging and arrhythmia. While short-term in late-life intervention of 10 mM NAM ameliorated age-associated cardiac arrhythmia, chronic or high-dose exposure led to detrimental cardiac effects, partly via impaired SERCA2a activity due to NAM-mediated inhibition of SIRT. This study highlights the narrow therapeutic window of NAM on cardioprotection with respect to both dose and duration, especially in aged populations with disrupted NAD⁺ metabolism. It also raises great cautions on the safety of supraphysiological NAM doses in both healthy and diseased individuals.

## Supporting information

Supplementary video1-100 mM NAM HL-1 cells

Supplementary video1-Control HL-1 cells

## References

1. Xie S, Xu S-C, Deng W, Tang Q. Metabolic landscape in cardiac aging: insights into molecular biology and therapeutic implications. Signal Transduction and Targeted Therapy. 2023;8:114. doi: 10.1038/s41392-023-01378-8

2. Wilkinson C, Todd O, Clegg A, Gale CP, Hall M. Management of atrial fibrillation for older people with frailty: a systematic review and meta-analysis. Age Ageing. 2019;48:196–203. doi: 10.1093/ageing/afy180

3. Li H, Hastings MH, Rhee J, Trager LE, Roh JD, Rosenzweig A. Targeting Age-Related Pathways in Heart Failure. Circulation Research. 2020;126:533–551. doi: 10.1161/CIRCRESAHA.119.315889

4. McReynolds MR, Chellappa K, Chiles E, Jankowski C, Shen Y, Chen L, Descamps HC, Mukherjee S, Bhat YR, Lingala SR, et al. NAD+ flux is maintained in aged mice despite lower tissue concentrations. Cell Systems. 2021;12:1160–1172.e1164. doi: 10.1016/j.cels.2021.09.001

5. Zhang X, Zhang Y, Sun A, Ge J. The effects of nicotinamide adenine dinucleotide in cardiovascular diseases: Molecular mechanisms, roles and therapeutic potential. Genes & Diseases. 2022;9:959–972. doi: 10.1016/j.gendis.2021.04.001

6. Norambuena-Soto I, Deng Y, Brenner C, Lavandero S, Wang ZV. NAD in pathological cardiac remodeling: Metabolic regulation and beyond. Biochim Biophys Acta Mol Basis Dis. 2024;1870:167038. doi: 10.1016/j.bbadis.2024.167038

7. Zhang D, Hu X, Li J, Liu J, Baks-te Bulte L, Wiersma M, Malik N-u-A, van Marion DMS, Tolouee M, Hoogstra-Berends F, et al. DNA damage-induced PARP1 activation confers cardiomyocyte dysfunction through NAD+ depletion in experimental atrial fibrillation. Nature Communications. 2019;10:1307. doi: 10.1038/s41467-019-09014-2

8. Lee CF, Chavez JD, Garcia-Menendez L, Choi Y, Roe ND, Chiao YA, Edgar JS, Goo YA, Goodlett DR, Bruce JE, et al. Normalization of NAD+ Redox Balance as a Therapy for Heart Failure. Circulation. 2016;134:883–894. doi: 10.1161/circulationaha.116.022495

9. Abdellatif M, Sedej S, Kroemer G. NAD^+^ Metabolism in Cardiac Health, Aging, and Disease. Circulation. 2021;144:1795–1817. doi: 10.1161/circulationaha.121.056589

10. Abdellatif M, Trummer-Herbst V, Koser F, Durand S, Adão R, Vasques-Nóvoa F, Freundt JK, Voglhuber J, Pricolo MR, Kasa M, et al. Nicotinamide for the treatment of heart failure with preserved ejection fraction. Sci Transl Med. 2021;13. doi: 10.1126/scitranslmed.abd7064

11. Zhang D, Wu CT, Qi X, Meijering RA, Hoogstra-Berends F, Tadevosyan A, Cubukcuoglu Deniz G, Durdu S, Akar AR, Sibon OC, et al. Activation of histone deacetylase-6 induces contractile dysfunction through derailment of α-tubulin proteostasis in experimental and human atrial fibrillation. Circulation. 2014;129:346–358. doi: 10.1161/circulationaha.113.005300

12. Qu W, Ralto KM, Qin T, Cheng Y, Zong W, Luo X, Perez-Pinzon M, Parikh SM, Ayata C. NAD+ precursor nutritional supplements sensitize the brain to future ischemic events. Journal of Cerebral Blood Flow & Metabolism. 2023;43:37–48. doi: 10.1177/0271678×231156500

13. Hoane MR, Pierce JL, Holland MA, Anderson GD. Nicotinamide treatment induces behavioral recovery when administered up to 4 hours following cortical contusion injury in the rat. Neuroscience. 2008;154:861–868. doi: 10.1016/j.neuroscience.2008.04.044

14. Hwang ES, Song SB. Possible Adverse Effects of High-Dose Nicotinamide: Mechanisms and Safety Assessment. Biomolecules. 2020;10. doi: 10.3390/biom10050687

15. Griffin SM, Pickard MR, Orme RP, Hawkins CP, Fricker RA. Nicotinamide promotes neuronal differentiation of mouse embryonic stem cells in vitro. NeuroReport. 2013;24:1041–1046. doi: 10.1097/wnr.0000000000000071

16. Li D, Tian Y-J, Guo J, Sun W-P, Lun Y-Z, Guo M, Luo N, Cao Y, Cao J-M, Gong X-J, et al. Nicotinamide supplementation induces detrimental metabolic and epigenetic changes in developing rats. British Journal of Nutrition. 2013;110:2156–2164. doi: 10.1017/S0007114513001815

17. Bitterman KJ, Anderson RM, Cohen HY, Latorre-Esteves M, Sinclair DA. Inhibition of Silencing and Accelerated Aging by Nicotinamide, a Putative Negative Regulator of Yeast Sir2 and Human SIRT1. Journal of Biological Chemistry. 2002;277:45099–45107. doi: 10.1074/jbc.M205670200

18. Carafa V, Rotili D, Forgione M, Cuomo F, Serretiello E, Hailu GS, Jarho E, Lahtela-Kakkonen M, Mai A, Altucci L. Sirtuin functions and modulation: from chemistry to the clinic. Clinical Epigenetics. 2016;8:61. doi: 10.1186/s13148-016-0224-3

19. Gorski PA, Jang SP, Jeong D, Lee A, Lee P, Oh JG, Chepurko V, Yang DK, Kwak TH, Eom SH, et al. Role of SIRT1 in Modulating Acetylation of the Sarco-Endoplasmic Reticulum Ca(2+)-ATPase in Heart Failure. Circ Res. 2019;124:e63–e80. doi: 10.1161/circresaha.118.313865

20. Roe AT, Frisk M, Louch WE. Targeting cardiomyocyte Ca2+ homeostasis in heart failure. Curr Pharm Des. 2015;21:431–448. doi: 10.2174/138161282104141204124129

21. Fink M, Callol-Massot C, Chu A, Ruiz-Lozano P, Izpisua Belmonte JC, Giles W, Bodmer R, Ocorr K. A new method for detection and quantification of heartbeat parameters in Drosophila, zebrafish, and embryonic mouse hearts. Biotechniques. 2009;46:101–113. doi: 10.2144/000113078

22. Cammarato A, Ocorr S, Ocorr K. Enhanced assessment of contractile dynamics in Drosophila hearts. Biotechniques. 2015;58:77–80. doi: 10.2144/000114255

23. Dobrev D, Wehrens XHT. Mouse Models of Cardiac Arrhythmias. Circ Res. 2018;123:332–334. doi: 10.1161/circresaha.118.313406

24. Nerbonne JM. Mouse models of arrhythmogenic cardiovascular disease: challenges and opportunities. Curr Opin Pharmacol. 2014;15:107–114. doi: 10.1016/j.coph.2014.02.003

25. Dautova Y, Zhang Y, Sabir I, Grace AA, Huang CL. Atrial arrhythmogenesis in wild-type and Scn5a+/delta murine hearts modelling LQT3 syndrome. Pflugers Arch. 2009;458:443–457. doi: 10.1007/s00424-008-0633-z

26. Soattin L, Lubberding AF, Bentzen BH, Christ T, Jespersen T. Inhibition of Adenosine Pathway Alters Atrial Electrophysiology and Prevents Atrial Fibrillation. Front Physiol. 2020;11:493–493. doi: 10.3389/fphys.2020.00493

27. Pillai VB, Samant S, Hund S, Gupta M, Gupta MP. The nuclear sirtuin SIRT6 protects the heart from developing aging-associated myocyte senescence and cardiac hypertrophy. Aging (Albany NY). 2021;13:12334–12358. doi: 10.18632/aging.203027

28. Covarrubias AJ, Perrone R, Grozio A, Verdin E. NAD+ metabolism and its roles in cellular processes during ageing. Nature Reviews Molecular Cell Biology. 2021;22:119–141. doi: 10.1038/s41580-020-00313-x

29. Ocorr K, Akasaka T, Bodmer R. Age-related cardiac disease model of Drosophila. Mech Ageing Dev. 2007;128:112–116. doi: 10.1016/j.mad.2006.11.023

30. Mouchiroud L, Houtkooper RH, Moullan N, Katsyuba E, Ryu D, Cantó C, Mottis A, Jo YS, Viswanathan M, Schoonjans K, et al. The NAD(+)/Sirtuin Pathway Modulates Longevity through Activation of Mitochondrial UPR and FOXO Signaling. Cell. 2013;154:430–441. doi: 10.1016/j.cell.2013.06.016

31. Jia H, Li X, Gao H, Feng Z, Li X, Zhao L, Jia X, Zhang H, Liu J. High doses of nicotinamide prevent oxidative mitochondrial dysfunction in a cellular model and improve motor deficit in a Drosophila model of Parkinson’s disease. J Neurosci Res. 2008;86:2083–2090. doi: 10.1002/jnr.21650

32. Khaidizar FD, Bessho Y, Nakahata Y. Nicotinamide Phosphoribosyltransferase as a Key Molecule of the Aging/Senescence Process. Int J Mol Sci. 2021;22. doi: 10.3390/ijms22073709

33. Wang MF, Hou C, Jia F, Zhong CH, Xue C, Li JJ. Aging-associated atrial fibrillation: A comprehensive review focusing on the potential mechanisms. Aging Cell. 2024;23:e14309. doi: 10.1111/acel.14309

34. Williams AC, Ramsden DB. Autotoxicity, methylation and a road to the prevention of Parkinson’s disease. Journal of Clinical Neuroscience. 2005;12:6–11. doi: 10.1016/j.jocn.2004.10.002

35. Liu L, Su X, Quinn WJ, 3rd, Hui S, Krukenberg K, Frederick DW, Redpath P, Zhan L, Chellappa K, White E, et al. Quantitative Analysis of NAD Synthesis-Breakdown Fluxes. Cell Metab. 2018;27:1067–1080.e1065. doi: 10.1016/j.cmet.2018.03.018

36. Mahmoud YI, Mahmoud AA. Role of nicotinamide (vitamin B3) in acetaminophen-induced changes in rat liver: Nicotinamide effect in acetaminophen-damged liver. Experimental and Toxicologic Pathology. 2016;68:345–354. doi: 10.1016/j.etp.2016.05.003

37. Shi W, Hegeman MA, Doncheva A, Bekkenkamp-Grovenstein M, de Boer VCJ, Keijer J. High Dose of Dietary Nicotinamide Riboside Induces Glucose Intolerance and White Adipose Tissue Dysfunction in Mice Fed a Mildly Obesogenic Diet. Nutrients. 2019;11. doi: 10.3390/nu11102439

38. Shi W, Hegeman MA, van Dartel DAM, Tang J, Suarez M, Swarts H, van der Hee B, Arola L, Keijer J. Effects of a wide range of dietary nicotinamide riboside (NR) concentrations on metabolic flexibility and white adipose tissue (WAT) of mice fed a mildly obesogenic diet. Mol Nutr Food Res. 2017;61. doi: 10.1002/mnfr.201600878

39. Perry CE, Halawani SM, Mukherjee S, Ngaba LV, Lieu M, Lee WD, Davis JG, Adzika GK, Bebenek AN, Bazianos DD, et al. NAD+ precursors prolong survival and improve cardiac phenotypes in a mouse model of Friedreich’s Ataxia. JCI Insight. 2024;9. doi: 10.1172/jci.insight.177152

40. van der Stelt I, Shi W, Bekkenkamp-Grovenstein M, Zapata-Pérez R, Houtkooper RH, de Boer VCJ, Hegeman MA, Keijer J. The female mouse is resistant to mild vitamin B3 deficiency. European Journal of Nutrition. 2021. doi: 10.1007/s00394-021-02651-8

41. Odening KE, Deiß S, Dilling-Boer D, Didenko M, Eriksson U, Nedios S, Ng FS, Roca Luque I, Sanchez Borque P, Vernooy K, et al. Mechanisms of sex differences in atrial fibrillation: role of hormones and differences in electrophysiology, structure, function, and remodelling. Europace. 2019;21:366–376. doi: 10.1093/europace/euy215

42. Siddiqi HK, Vinayagamoorthy M, Gencer B, Ng C, Pester J, Cook NR, Lee IM, Buring J, Manson JE, Albert CM. Sex Differences in Atrial Fibrillation Risk: The VITAL Rhythm Study. JAMA Cardiol. 2022;7:1027–1035. doi: 10.1001/jamacardio.2022.2825

