## Supplementary files for "Dual effects of nicotinamide on aging-related arrhythmia: protective at low dose, proarrhythmic at high doses"

**Supplementary information**

**Table S1. Lifespan parameters of *Drosophila melanogaster* supplemented with different doses of nicotinamide (NAM)**

| | Groups | Control | 10 mM NAM | 20 mM NAM | 50 mM NAM | 100 mM NAM |
|---|---|---|---|---|---|---|
| Male | Median survival | 67 | 69 | 58 | 28 | 3 |
| | 90% mortality | 79 | 77 | 65 | 39 | 7 |
| | Average lifespan | 63.2 | 55.3 | 48.8 | 24.5 | 3.7 |
| | Log-rank test* (over the Control) | | 0.8831 | <0.0001 | <0.0001 | <0.0001 |
| | Hazard ratios** (95% CI of ratio over the Control) | | 1.022 (0.752 - 1.389) | 2.783 (1.972 - 3.929) | 4.620 (3.083 - 6.922) | 4.527 (2.936 - 6.981) |
| Female | Median survival | 68 | 79 | 57 | 34 | 4 |
| | Maximum survival | 84 | 87 | 77 | 51 | 5 |
| | Average lifespan | 60.4 | 67.2 | 48.0 | 31.7 | 3.8 |
| | Log-rank test* (over the Control) | | 0.0006 | 0.0002 | <0.0001 | <0.0001 |
| | Hazard ratios** (95% CI of ratio over the Control) | | 0.6039 (0.441- 0.827) | 1.74 (1.254 - 2.416) | 3.455 (2.388 - 5.000) | 3.956 (2.686 - 5.826) |

Median survival, age of 90% mortality, and average lifespan are in days.

*Log-rank test and **hazard ratios were calculated relative to the control group.

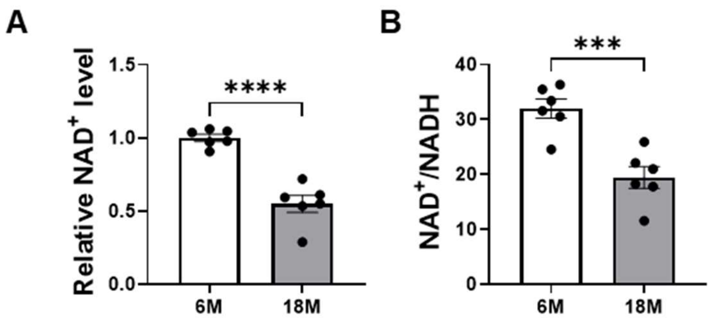


**Figure S1. NAD$^+$ and NAD$^+$/NADH levels in mouse atrial heart tissue.** (A) Relative NAD$^+$ levels and (B) NAD$^+$/NADH ratio of atrial tissue from 6-month old (6M) and 18-month old (18M) mice (n=6, respectively). Data are expressed as mean ± standard error of the mean. Two tailed unpaired Student t-test was used. ***$P < 0.001$, ****$P < 0.0001$.

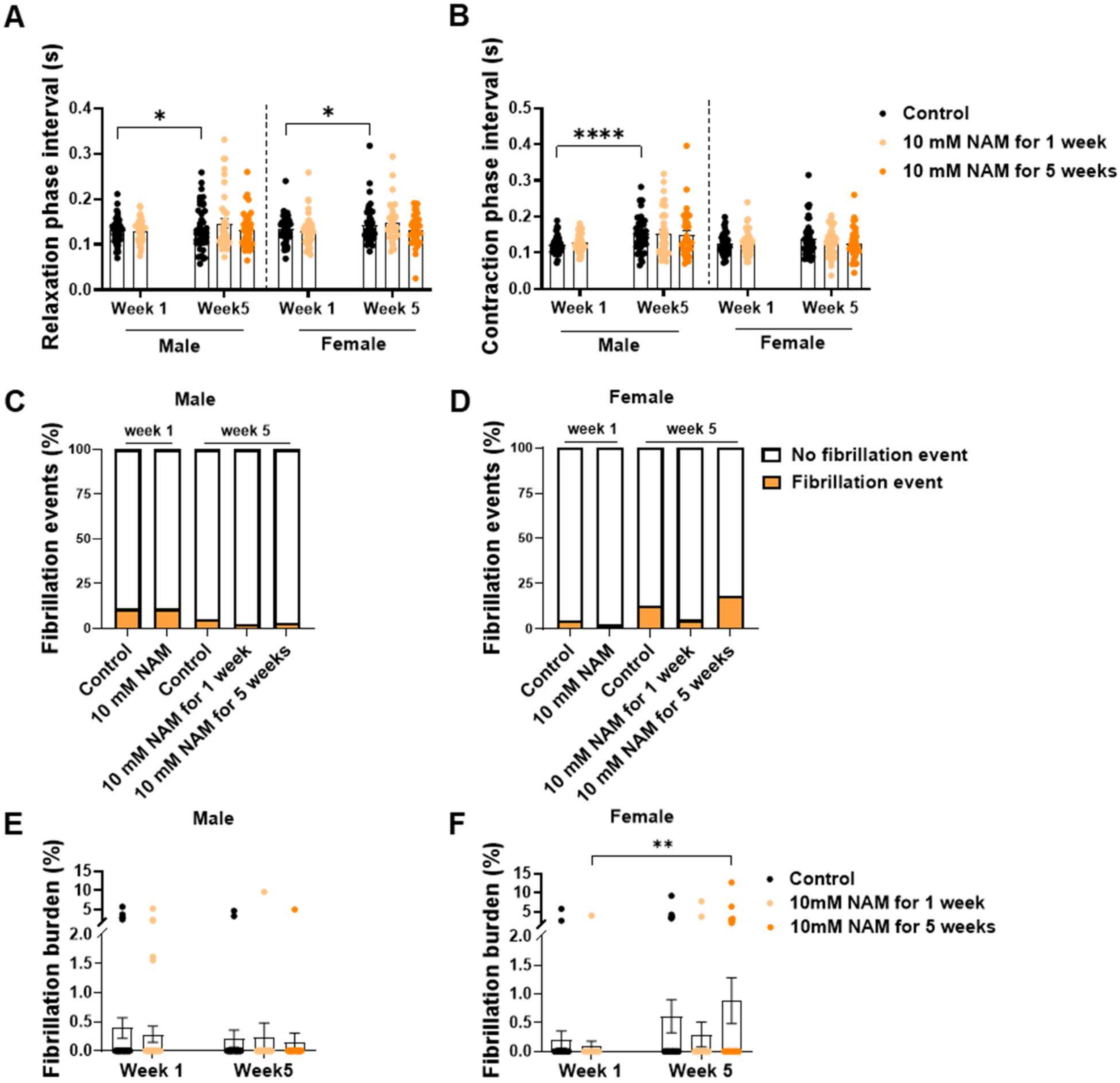


**Figure S2. Cardiac contractile function and fibrillation events and burden of *Drosophila melanogaster* treated with 10 mM nicotinamide (NAM) compared to 0 mM NAM (control).** (A) Relaxation phase intervals, (B) contraction phase intervals, (C&D) fibrillation event incidence, and (E&F) fibrillation burden of male and female 1-week-old flies fed with a control diet (n=45 males, 41 females) or a 10 mM NAM-supplemented diet (n=45 males, 44 females), and 5-week-old flies fed with a control diet (n=38 males, 39 females), a 10 mM NAM-supplemented diet for 1 week (n=40 males, 39 females), or for 5 weeks (n=33 males, 37 females). Data are expressed as mean ± standard error of the mean. Chi-square test was used for fibrillation events and Scheirer-

Ray-Hare test followed by Dunn's post-hoc test was used for fibrillation burden. *$P < 0.05$, ** $P < 0.01$.

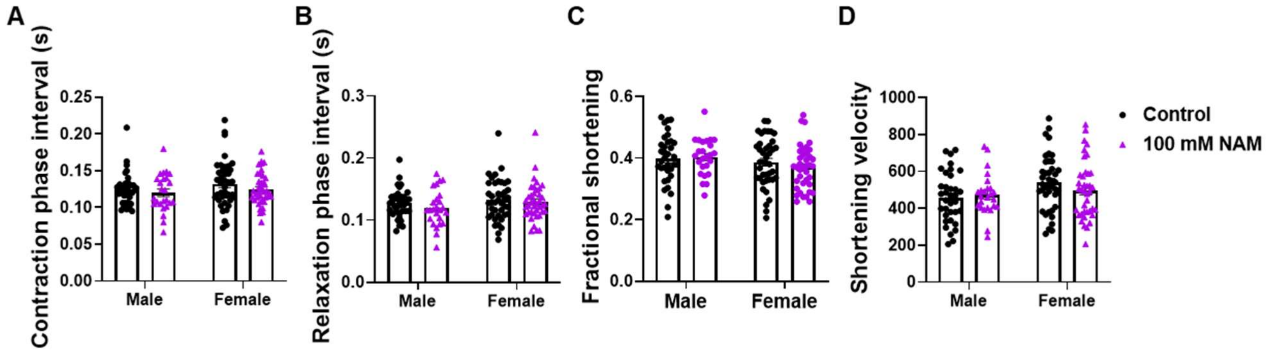


**Figure S3. Cardiac contractile function of *Drosophila melanogaster* treated with 100 mM nicotinamide (NAM) compared to 0 mM NAM (control).** (A) Contraction phase intervals, (B) relaxation phase intervals, (C) fractional shortening, and (D) shortening velocity of semi-intact heart preparations of the control (n=35 males, 41 females) and 100 mM NAM-treated flies (n=25 males, 39 females). Data are expressed as mean ± standard error of the mean. Two-way ANOVA followed by post-hoc Fisher's LSD test was used.

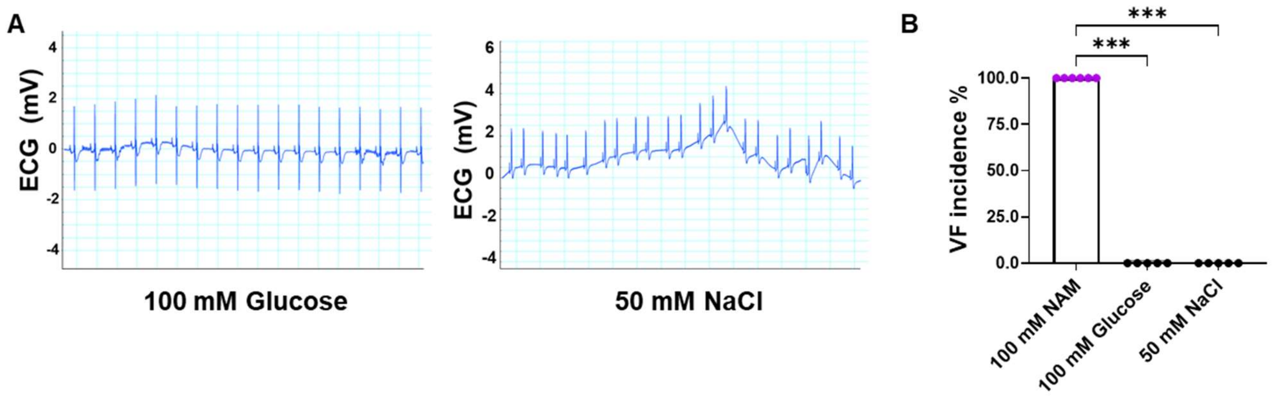


**Figure S4. Electrogram (ECG) recording of *ex vivo* mouse hearts perfused with high-dose glucose and NaCl.** (A) Representative ECG traces of *ex vivo* mouse hearts perfused with an equivalent osmotic molar concentration of 100 mM glucose or 50 mM NaCl (n=5 males, respectively). (B) Incidence of ventricular fibrillation (VF) in the *ex vivo* mouse hearts perfused with either 100 mM NAM, 100 mM glucose, or 50 mM NaCl. High-dose glucose and NaCl did not induce VF, in contrast to 100 mM nicotinamide (NAM, n=4 males, 2 females). Kruskal-Wallis test followed by Dunn's post-hoc test was used. ***$P < 0.001$.
